# The functional impact of 1,570 SNP-accessible missense variants in human *OTC*

**DOI:** 10.1101/2022.10.26.513893

**Authors:** Russell S. Lo, Gareth A. Cromie, Michelle Tang, Kevin Teng, Katherine Owens, Amy Sirr, J. Nathan Kutz, Richard N. McLaughlin, Hiroki Morizono, Ljubica Caldovic, Nicholas Ah Mew, Andrea Gropman, Aimée M. Dudley

## Abstract

Deleterious mutations in the X-linked gene encoding ornithine transcarbamylase (*OTC*) cause the most common urea cycle disorder, OTC deficiency. This rare, but highly actionable disease can present with severe neonatal onset in males or with later onset in either sex. Neonatal onset patients appear normal at birth but rapidly develop hyperammonemia, which can progress to cerebral edema, coma and death, outcomes ameliorated by rapid diagnosis and treatment. Existing biochemical assays have limitations, including the sensitivity of the citrulline assays used in newborn screening panels. With prior knowledge of variant pathogenicity, DNA sequence-based diagnostics would provide an alternative screening method. Here, we develop a high throughput functional assay for human OTC and measure the impact of 1,570 variants, 84% of all SNP-accessible missense mutations. Our assay scores agree well with existing clinical significance calls, distinguishing known benign from pathogenic variants and variants with neonatal onset from late-onset disease presentation. Further, use of an intronless expression construct allows us to measure the impact of amino acid changes at splice sites independent of their effect on splicing, thereby separating the contribution of splicing and protein coding changes to aid the analysis of molecular mechanisms underlying pathogenicity. Finally, we assess the utility of our functional data on *OTC* variant curation by using the current ACMG/AMP guidelines to reclassify variants. Inclusion of our data as PS3/BS3 substantially improves variant interpretation. Thus, our dataset is of high clinical utility and illustrates the power of functional assays to inform interpretation of existing and novel genetic variation.

## Introduction

Urea cycle disorders (UCD) are Inborn Errors of Metabolism caused by deficiencies in any of the eight proteins in the pathway responsible for the conversion of nitrogen, a waste product of protein metabolism, to urea. Accumulation of nitrogen, in the form of ammonia, to toxic levels in the blood and brain leads to symptoms that include vomiting, lethargy, behavioral abnormalities, cerebral edema, seizures, coma and death. Ornithine transcarbamylase (OTC) deficiency is the most common UCD, accounting for approximately half of all diagnosed cases.^1^ OTC is a nuclear-encoded mitochondrial protein that converts ornithine and carbamoyl phosphate to citrulline in human liver cells. As an X-linked trait, severe disease more often presents in males, but can also occur in females harboring pathogenic alleles. The disease has both strong genetic and environmental components.^2^ While severe loss of function mutations are often associated with neonatal presentation, age of onset can be variable, even among patients harboring the same *OTC* allele.^3–5^ Late-onset presentation is associated with hyperammonemia-triggering events, including high protein intake, prolonged fasting, surgery, exposure to organic chemicals, pregnancy and administration of specific medications, such as corticosteroids or valproate.^6^ Severe outcomes from these events can be prevented by rapid diagnosis and administration of nitrogen scavengers.^7^ In patients presenting in the neonatal period, liver transplantation is an effective method of preventing further hyperammonemia events.^8^

Unfortunately, in the absence of family history, presymptomatic diagnosis of OTC deficiency is challenging. The sensitivity and stability of the metabolite-based assay used for newborn screening of OTC deficiency is limited, and currently this disorder is screened in only eight U.S. states.^9^ In severely affected males, hyperammonemia can occur rapidly after birth, often before newborn screening results are available. Furthermore, in females, the heterogeneity of liver cells due to X-inactivation confounds enzymatic assays from liver biopsies. Thus, DNA sequencing-based diagnostics have the potential to reduce the morbidity and mortality associated with delayed treatment or underdiagnosis.

Because this devastating disease is highly actionable, *OTC* was added to the American College of Medical Genetics and Genomics secondary findings gene list (ACMG SF v2.0)^10^ which offers guidance for which genes should be evaluated when an individual undergoes clinical exome and genome sequencing for any reason. Inclusion of *OTC* on this small list of high priority genes highlights the utility of genetic testing. However, DNA sequencing often returns novel sequence variants of uncertain significance (VUS). Approximately 20-30% of OTC deficiencies occur *de novo*,^11^ and only a minority of OTC variants are recurrent.

Advances in the cost and scale of both DNA sequencing and synthesis have enabled the development of high throughput functional assays to experimentally address this problem. These so-called deep mutational scanning (DMS) or multiplexed assays for variant effect (MAVEs) approaches have been applied to the comprehensive analysis of amino acid or nucleotide variants in synthetic peptides, protein domains, whole proteins, transcriptional promoters, RNA transcripts, splice sites, and DNA replication origins.^12–14^ Many of these studies leverage the experimental tractability of model organisms to perform low cost, high throughput studies on massively parallel scales. Because fundamental biological processes are often highly conserved between humans and model organisms, many human protein coding sequences are able to functionally replace their orthologs in model organisms such as the yeast, *Saccharomyces cerevisiae*.^15–23^ This is especially true for genes that encode core metabolic processes such as nutrient biosynthesis and utilization.^22^

The enzymes in the urea cycle responsible for arginine biosynthesis are highly conserved between humans and yeast (Figure 1). At the protein level, human OTC is 34.5% identical and 54.5% similar to its yeast ortholog, Arg3. Here, we present a complementation assay in which growth of yeast cells in the absence of arginine serves as a proxy for the function of human OTC. We use this assay at scale to measure relative growth scores for 1,570 SNP-accessible single amino acid substitutions across the length of the protein. Our results, based on multiple biological and experimental replicates of sequence-confirmed clones, are highly reproducible. A large proportion of these missense variants display little to no protein function, suggesting that OTC is highly sensitive to amino acid substitutions. Variants within this extremely low range of activity include the majority of amino acid substitutions in the active sites of the enzyme, variants resulting from SNPs with pathogenic or likely-pathogenic clinical classifications, and variants resulting from SNPs associated with severe disease (neonatal onset) in the literature. Finally, we highlight examples of the utility of our functional data for SNP interpretation in which its use as PS3/BS3 evidence enables the reclassification of VUS.

**Figure 1.**
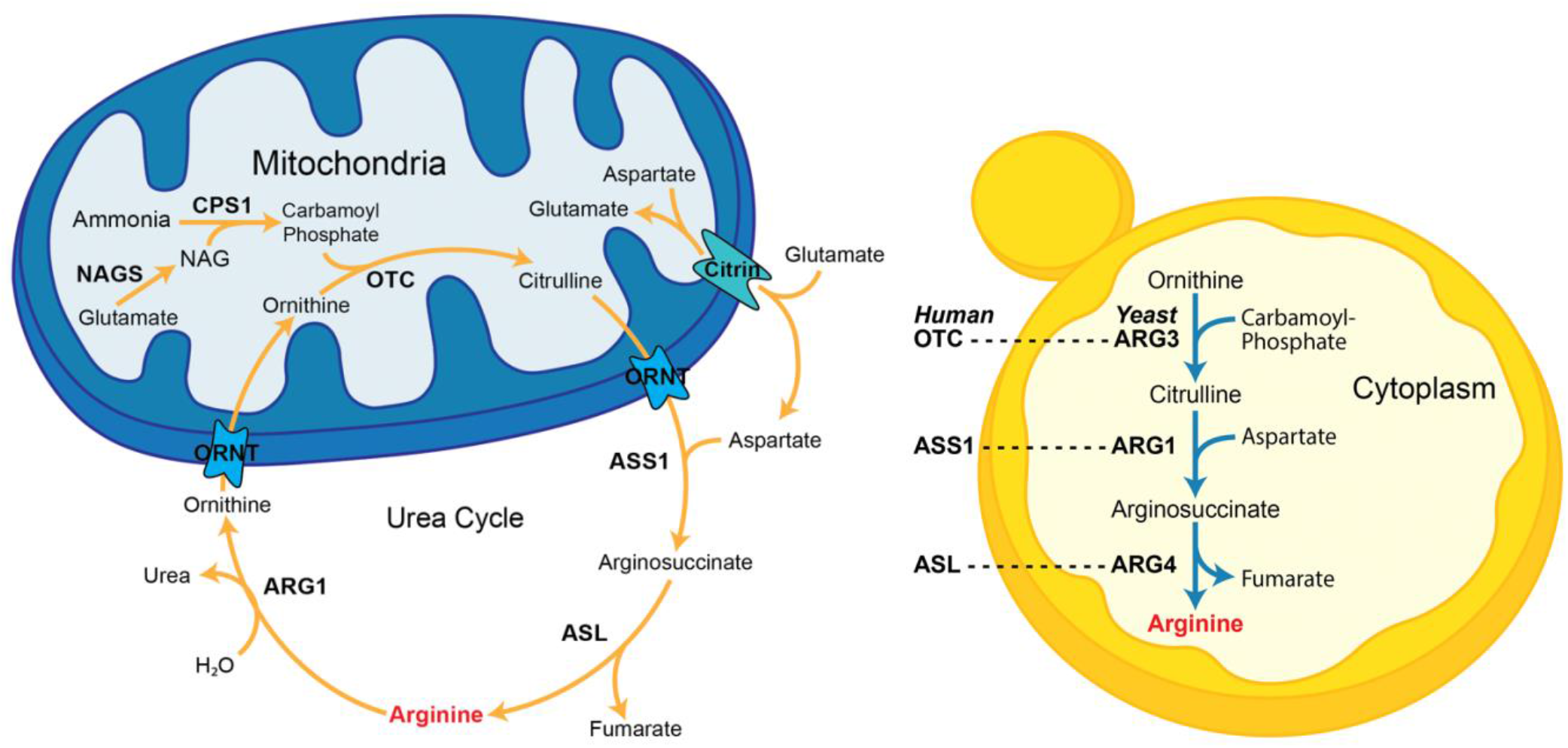
Comparison of the urea cycle in humans and the arginine biosynthesis pathway in *Saccharomyces cerevisiae*. Shown in blue is a human liver mitochondrion which harbors the enzymes N-acetylglutamate synthase (*NAGS*), carbamylphosphate synthetase 1 (*CPS1*), ornithine transcarbamylase (*OTC*), ornithine transporter (*ORNT*) and citrin transporters. Carbamoyl phosphate and ornithine are bound by ornithine transcarbamylase (*OTC*) and converted into citrulline. Citrulline and aspartate are then converted to arginine via the enzymatic activity of arginosuccinate synthase I (*ASS1*) and arginosuccinate lyase (*ASL*). Shown in yellow is a yeast cell containing the orthologous enzymes ornithine carbamoyltransferase (*ARG3*), arginosuccinate synthetase (*ARG1*), and argininosuccinate lyase (*ARG4*) which act in the cytoplasm to convert ornithine, carbamoyl phosphate, and aspartate to arginine.

## Methods

### Plasmid and strain construction

All *Saccharomyces cerevisiae* strains used in this study (Supplemental Table S1) were derived from the isogenic lab strain FY4.^24^ Unless noted, strains were grown in rich YPD media (1% yeast extract, 2% peptone, and 2% glucose) or minimal (SD) medium (without amino acids, 2% glucose) using standard media conditions and methods for yeast genetic manipulation.^25^

To construct yeast strains harboring OTC variants, the *ARG3* open reading frame was first deleted from FY4 and replaced with a selectable kanamycin resistance gene from pFA6a-kanMX6^26^. In this strain, expression of the kanamycin resistance gene was under control of the *S. cerevisiae ARG3* promoter and the *A. gossypii* TEF terminator.

The OTC protein coding sequence (GenBank: NM_000531.6) was optimized for expression in yeast (hereafter *yOTC*, GenBank: ON872185), using a custom, in-house method designed to match the codon usage frequency of *S. cerevisiae*. Briefly, at amino acid positions conserved between the yeast and human proteins, the yeast (S288c reference) codon was used. In non-conserved locations, codons encoding the appropriate OTC amino acid were chosen to be as similar as possible to the usage frequency of the codons at corresponding positions in *ARG3*.

The mitochondrial leader sequence (amino acids 2-32) of the human OTC was omitted from *yOTC*, and *yOTC* was placed under the control of the native *ARG3* promoter. However, from hereon, stated variant amino acid positions refer to positions of the native, full-length OTC protein. To avoid disrupting the regulation of the neighboring yeast gene (*TRL1*), which shares a 186 bp terminator sequence with *ARG3*, we retained the *TRL1* terminator and introduced a *TRP2* terminator sequence upstream of the drug marker to control the termination of yOTC (Figure 2).

**Figure 2.**
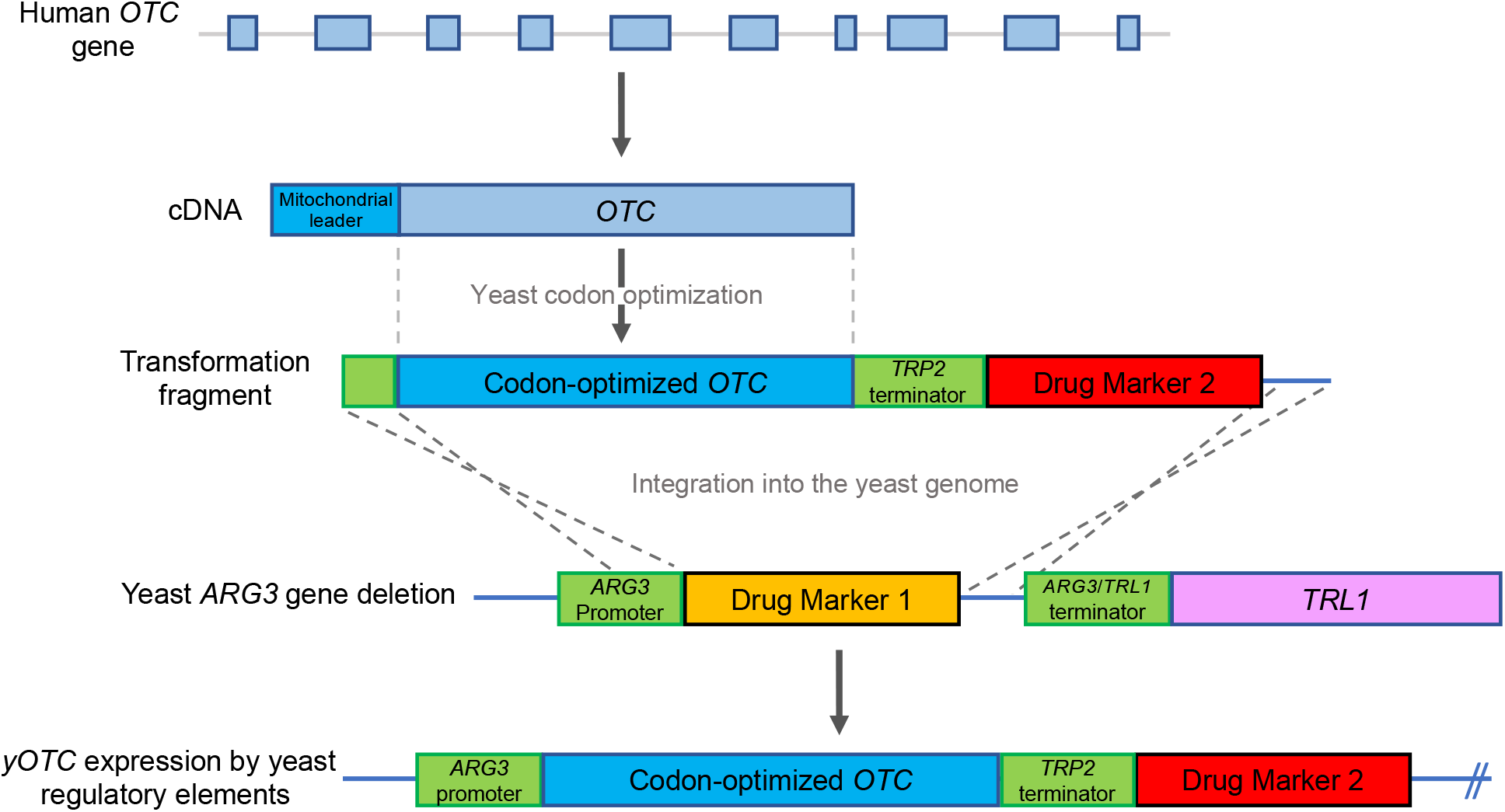
Experimental design. Starting with a cDNA sequence of human *OTC*, the mitochondrial leader sequence was omitted and the remaining sequence was codon optimized for improved expression in yeast. For strain construction, the *ARG3* ORF was deleted in a haploid FY4 laboratory yeast strain and replaced with drug marker 1. This null deletion strain was then transformed with PCR products containing either wild type or variant alleles of the yeast codon-optimized human *OTC* protein coding sequence (*yOTC*) along with drug marker 2. The resulting transformants’ expression of *yOTC* is under the control of yeast regulatory elements (ARG3 promoter and TRP2 terminator).

A plasmid containing the wild type *yOTC* was constructed as follows. A gBLOCK containing the *ARG3* promoter (*pARG3*) and *yOTC* was synthesized (Genewiz) and cloned upstream of the *TRP2* terminator and NatMX6 drug marker cassette in a pUC19-based vector using the NEB HiFi Assembly kit (New England Biolabs). This AB614_yOTC plasmid (GenBank: ON872185) was used as a template for variant library construction.

The yOTC variant library, designed to include all 1,879 SNP-accessible amino acid substitutions (excepting the start and stop codons), was synthesized by Twist Bioscience. Each variant fragment synthesized consists of a *pARG3-yOTC-tTRP2-NatMX* cassette from the AB614_yOTC template plasmid and each amino acid substitution was encoded by a single codon (Supplementary Table S2). In addition, 19 bp of additional *tTEF* homology (5’ TCCTTCTTTCGTGTTCTTA 3’) was added to the cassette during library synthesis to the 3’ end. Primers corresponding to 16 bp of *pARG3* homology (5’ GAGCCGGATTGGTCAC 3’) and the 19 bp of additional *tTEF* homology (5’ TCCTTCTTTCGTGTTCTTA 3’) were used to prime a 15-cycle PCR amplification specific for the Twist variant fragment versus the original wildtype *yOTC* vector template.

Approximately 500 ng of each PCR-amplified variant fragment was transformed into the *arg3*Δ*2∷KAN-tTEF* strain using standard methods (Figure 2). 5700 individual transformants, each harboring a different variant, were picked into 96-well plates containing rich media supplemented with 100 μg/mL nourseothricin. Each library plate also contained replicates of the same control strains: *arg3*Δ*0∷NATMX* deletion (n = 2 wells) and wild type *yOTC* (n = 4 wells) for use in phenotype normalization.

### Determining the Variant Encoded by Each Transformation Isolate

Transformations were carried out using DNA pools, each consisting of all variants at a particular codon. Therefore, for each isolate we know the target codon, but not which specific variant of that codon is present. To determine the specific variant in each isolate, and to filter out clones harboring secondary mutations, we developed an Oxford Nanopore MinION sequencing pipeline and confirmed the findings using Illumina sequencing (Supplementary File 1).

Briefly, we pooled and sequenced multiple isolates, each with a different target codon. Reads from each pool were then aligned to the reference *yOTC* sequence. At each of the target codons in the pool, we then determined the most frequent variant codon. If this codon passed quality control, it was accepted as the variant at that target codon. As we know which isolate in the pool had which target codon, this allowed us to determine the variant codon in each isolate. Any secondary mutations occurring in each isolate were also recorded.

To validate our Oxford Nanopore pipeline, the sequence at the target codon in each isolate was also determined by Illumina sequencing (Supplementary File 1). For the variant calls passing quality control in the MinIon pipeline, 99.8% agreed with the Illumina sequencing variant call.

### Measuring the Growth of Each Isolate

The growth of each isolate was estimated as described in Supplementary File 1. Briefly, each isolate was grown in 96-well format in rich liquid medium (containing arginine) until saturated. Each isolate was then pinned robotically from the non-selective liquid medium to selective solid minimal medium plates (lacking arginine). Plates were grown at 30° C for 72 hours and imaged at each 24-hour interval. Growth for day 3 images was quantified by generating a pseudo patch “volume” for each pinned isolate consisting of the product of the patch area and the mean patch pixel intensity (grayscale).

### Data Normalization and Estimating the Relative Growth of Each Variant Genotype

Raw phenotypic values were normalized, quality control filters were applied to each isolate, and a final relative growth estimate for each genotype was determined, as described in Supplementary File 1. Briefly, normalization steps to account for the effects of plate-to-plate variation, relative growth of neighboring patches, and plate edge effects were applied to the data using a custom script. Isolates with (non-synonymous) secondary mutations or outlier phenotypes were removed from the dataset. Finally, a linear model was used to estimate the relative growth of each genotype, on a scale with growth of null controls set to 0 and growth of wild type *yOTC* set to 1.

### Initial Assay with Known Pathogenic and Benign Variants

For our initial assay (Figure 3), we tested the common K46R benign variant and the known R141Q null pathogenic variant against the *arg3* deletion strain and the wild type yOTC variant. Pinning assays were performed in checkerboard format with multiple isolates using internal wells/spots such that no data normalization was needed to adjust for plate edge effects and neighbor growth effects. Images were processed as described above and mean growth values relative to wild type were calculated with null subtraction.

**Figure 3.**
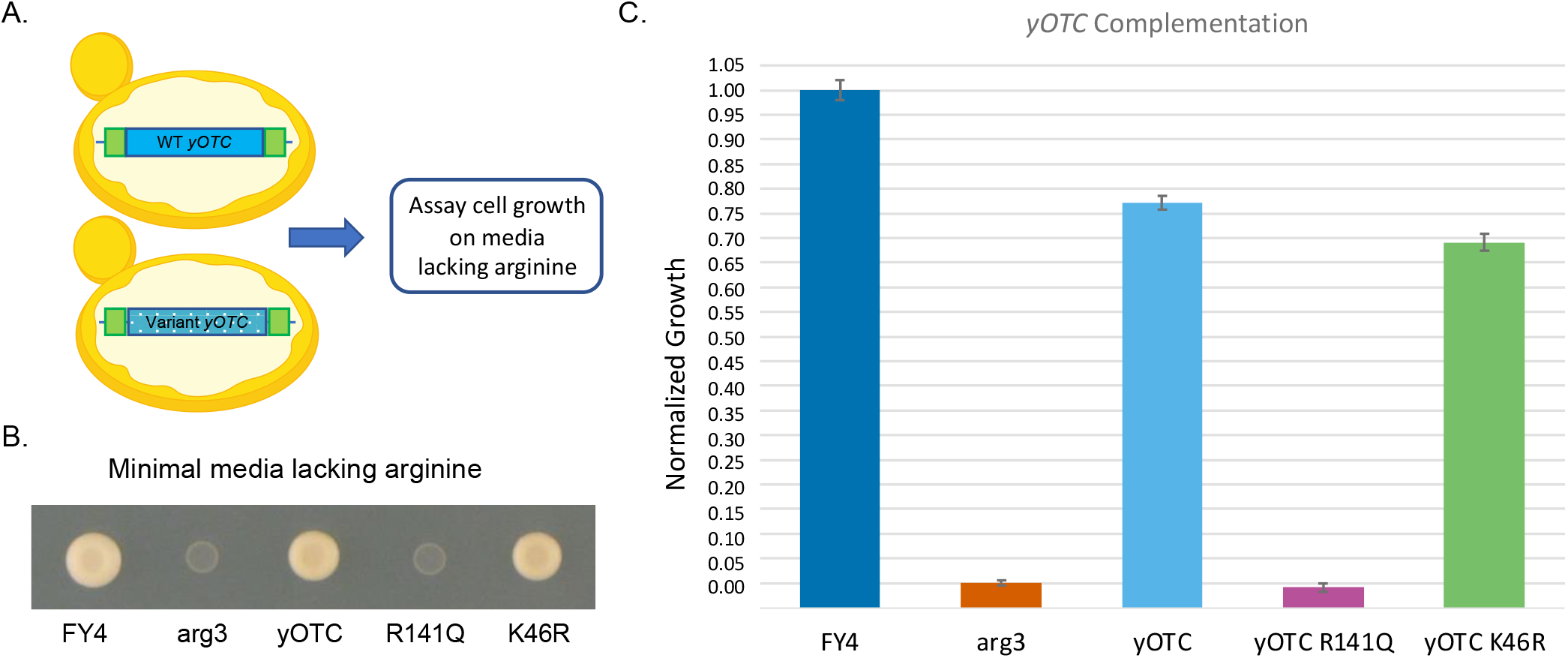
Establishing a yeast model for human OTC function. A) Haploid yeast strains harboring single integrated copies of wild type or variant *yOTC* can be assayed for growth on minimal media lacking arginine. B) Yeast pinning assay which demonstrates that yOTC wild type and the K46R benign variant can complement the arg3 deletion. Growth on minimal media after 3 days is shown. C) Wild type *yOTC* can complement the yeast *ARG3* deletion by 77% as assayed by the quantitative growth of yeast cells (product of the area of growth and the intensity) pinned onto minimal medium lacking arginine. Growth estimates for each genotype ± their standard deviations are displayed relative to wild type FY4 yeast which is set to 1.0 and the *arg3* deletion strain which is set to 0. Assays were performed in replicate (4 independent FY4 clones, 2 *arg3D0∷KANMX* clones, 4 WT *yOTC* clones, 2 independent R141Q clones, and 6 independent K46R clones with 3 technical replicates of pinning plates).

### ClinVar Classification and Clinical Stratification

For ClinVar and clinical stratification, we assign a classification to an amino acid substitution if any SNP encoding that substitution has the classification.

### Empirical P-Values

For calculations of empirical P-values, 100k samples of the appropriate size were drawn, without replacement, from the appropriate subset of assay results. These samples were used to calculate a null distribution of medians, or differences in medians, to which the test median or difference in medians was compared.

## Results

### Human OTC can functionally replace ARG3 in yeast

Our strategy for optimizing yeast complementation assays is based on the guiding principle that complementation by a human protein coding sequence is best achieved by closely matching the subcellular localization and expression of the yeast ortholog. The X-linked *OTC* gene contains ten exons and nine introns. Because, like most genes in the yeast genome, the yeast *OTC* ortholog (*ARG3*) is intronless,^27^ we chose the OTC cDNA sequence (GenBank: NM_000531.6). The 354 amino acid OTC precursor protein contains a 32-amino acid mitochondrial leader sequence that is cleaved from the N-terminus following import.^28^ Since the Arg3 enzyme functions in the yeast cytosol,^29^ we omitted amino acids 2-32 from the human *OTC* coding sequence. However, to simplify comparisons with clinical literature and databases, variant amino acid positions within this protein are referred to using their position in the native, full-length OTC protein. Because OTC functions as a homotrimer with active sites formed at the monomer interfaces,^30^ we expected that additional human proteins would not be required for OTC enzyme function in yeast.

We next sought to match the gene expression of *OTC* to that of its yeast ortholog as follows (Figure 2). First, we generated a codon-harmonized version of *OTC* that encodes the human protein but more closely matches the codon utilization frequency of *ARG3* than the original OTC cDNA sequence (Materials and Methods). A single copy of this new sequence, hereafter called *yOTC*, was integrated into the yeast genome at the *ARG3* locus, under the control of the native *ARG3* promoter. This experimental design ensured stable maintenance of *yOTC* (wildtype and variant library) at a uniform copy number in all cells and placed *yOTC* under the native regulatory environment of the yeast ortholog.

### Establishing an in vivo OTC functional assay in yeast

Because ornithine transcarbamylase activity is required for yeast to grow in the absence of exogenously supplied arginine, growth on minimal medium (lacking arginine) is a quantitative measurement of enzyme function that is amenable to high throughput analysis. To quantify the ability of *yOTC* to functionally replace *ARG3*, we compared growth of the parental strain (FY4, expressing yeast *ARG3*), an *arg3* deletion strain (*arg3Δ0∷NATMX*, lacking any ornithine transcarbamylase), and a strain harboring *yOTC* at the *ARG3* locus as the sole source of enzyme activity. Each strain was grown to saturation under a nonselective condition (rich medium containing arginine), replica pinned to minimal medium lacking arginine, and allowed to grow at 30° C for 72 hours. As expected, FY4 grew robustly while the *arg3* deletion was unable to grow, displaying only the faint patch of cells deposited by the initial replica pinning from rich medium (Figure 3). In this assay, *yOTC* exhibited robust complementation of the *arg3* deletion, with 77% colony growth relative to the FY4 strain harboring the yeast ortholog (Figure 3).

To further test how well activity in yeast agreed with activity in humans, we constructed and assayed a small set of well characterized variants with pathogenic or benign clinical significance calls in ClinVar.^31^ We expected that amino acid changes corresponding to those of known pathogenic missense variants would impair OTC activity, resulting in poor growth, potentially down to the complete lack of growth conferred by the *arg3* deletion. In contrast, amino acid changes corresponding to those of benign missense variants should result in high growth, potentially up to the high level conferred by wildtype yOTC. The pathogenic 422G>A missense variant encodes the R141Q amino acid substitution that abolishes enzymatic activity without altering protein abundance.^32^ In our assay, the strain harboring this amino acid variant behaved like the *arg3* deletion mutant and failed to grow on minimal medium (Figure 3). Conversely, the common 119G>A benign missense variant encodes the K46R amino acid substitution, which in our assay functionally complemented the *arg3* deletion to 90% growth relative to wildtype yOTC (Figure 3). Thus, *yOTC* complements *ARG3*, and the activity of amino acid substitutions corresponding to known pathogenic and benign missense variants are consistent with expectation.

### Assaying a Library of SNP-accessible OTC Amino Acid Substitutions

With a validated functional assay in hand, we proceeded with the construction and analysis of a full-scale library. We chose to comprehensively analyze the effect of single amino acid substitutions that are accessible by a single nucleotide polymorphism (SNP), as these are variants that are likely to arise in the human population. Similar to enzymatic assays, our analysis does not directly test individual SNPs, rather it measures the impact of the amino acid substitutions encoded by the SNPs.

Yeast cells were transformed with a library of *yOTC* derivatives, each encoding a single SNP-accessible amino acid substitution (Supplemental Table S2). For each transformant, the full *yOTC* ORF was then sequenced to identify the codon change present using Oxford Nanopore and Illumina sequencing (Materials and Methods). For Oxford Nanopore data passing quality control, there was 99.8% agreement between the altered codon identified and that identified by Illumina sequencing (Materials and Methods). The Oxford Nanopore pipeline also identified transformants with indels, nonsynonymous, or nonsense secondary mutations, which were removed from the final dataset (Materials and Methods). For the majority of altered codons, multiple independent isolates were assayed. Together, we were able to isolate and assay 84% (1,570 out of 1,878) of the total possible SNP-accessible amino acid substitutions in *OTC* (Supplemental Table S1).

We next assayed the growth of each isolate individually on solid medium in a gridded array format. Strains were first grown to saturation in rich liquid medium (containing arginine) in 96-well plates. The liquid cultures were then replica pinned robotically to minimal medium lacking arginine and grown for 72 hours. Growth was measured as the product of the area and average pixel intensity of each patch, using a custom script (Materials and Methods). After normalization to account for plate-to-plate, edge, and neighbor patch effects on growth, a linear model was used to estimate growth of each yeast *yOTC* genotype (and single amino acid change) relative to wildtype *yOTC* (Materials and Methods). Wildtype *yOTC* growth was set to a value of 100%, with the *arg3* deletion strain set to a value of 0% (Supplemental Table S3).

In contrast to fitness-based, pooled approaches to phenotyping, individual phenotyping in this manner provides a stable estimate of activity for each *yOTC* genotype. Such measurements are independent of the composition of a particular competitive pool and not subject to relative fitness changes as the mean pool fitness increases over time. ^33^ This approach also facilitates the integration of the current dataset with future datasets assayed with a defined set of controls, allowing the additional genotypes to be added to the same scale and compared directly.

### A High Proportion of Amino Acid Substitutions Impair OTC Activity

The amino acid substitutions tested conferred a wide range of growth values spanning the range between the wild-type (*yOTC*) and null (*arg3*) controls (Figure 4). Strikingly, 27% of amino acid substitutions exhibit relative growth values below 5% of wild-type activity. The approximately normal distribution of these values centered on the null control is consistent with complete loss of OTC function and a small amount of measurement noise. The remaining amino acid substitutions exhibit a relatively uniform distribution across the rest of the growth range. Interestingly, there is no concentration of amino acid substitutions clustered around the wildtype level of growth and only 10% of amino acid substitutions (n= 162) had growth estimates equal to or exceeding that of wildtype yOTC. Because the human protein coding sequence did not fully complement loss of the yeast ortholog (77% of *ARG3* growth, Figure 3), the assay has the potential to detect activity-enhancing variants up to at least *ARG3* levels of growth. Interestingly, among the 1% of amino acid substitutions with the highest growth, 10/16 had amino acid substitutions at positions 143, 324-326, or 349-350. Visualizing activity in our assay versus the amino acid position shows a relatively uniform distribution across the length of the protein, with most positions exhibiting high levels of intolerance to amino acid substitution (Figure 5). Two small regions of the protein, one between amino acids 128-133 and another between amino acids 265-280, show higher tolerance with most variants having >50% activity (Figure 5). The region between residues 265 and 280 contains the conserved SMG loop, which may undergo a conformational change upon the binding of carbamoyl phosphate and ornithine that shields the active site of the enzyme.^34^ It has been suggested that despite its conservation, most amino acid changes in the loop would not necessarily affect its function^35^ and our results are consistent with this hypothesis.

**Figure 4.**
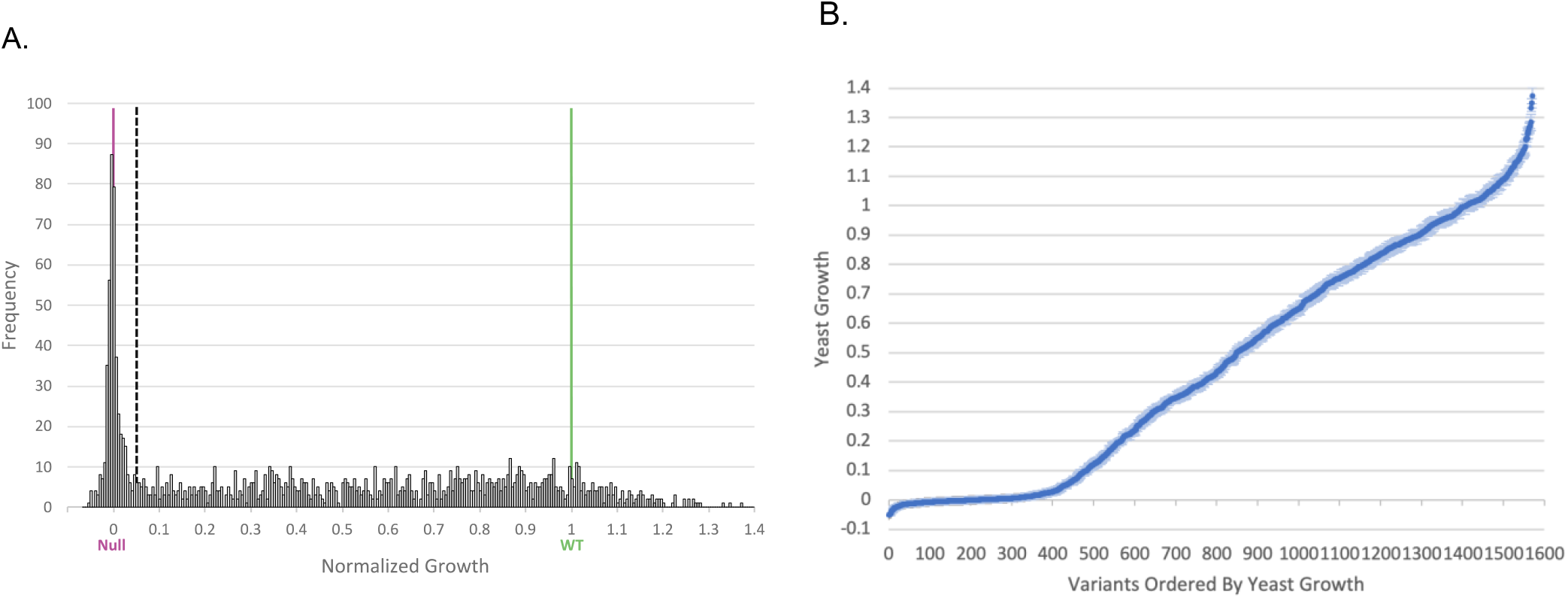
Quantitative growth measurements of 1,570 OTC variants. A) Frequency of yOTC variants exhibiting varying amounts of normalized growth. Levels of yOTC variant growth are reported relative to growth of the *arg3* deletion (normalized growth value = 0, marked w/ a purple line) and the wild type *yOTC* (normalized growth value = 1, marked with a green line). A 0.05 cutoff for high confidence deleterious enzyme activity is marked with a black dashed line. B) Relative growth of each yOTC variant ordered on the x-axis and plotted with +/− SE of the growth estimates.

**Figure 5.**
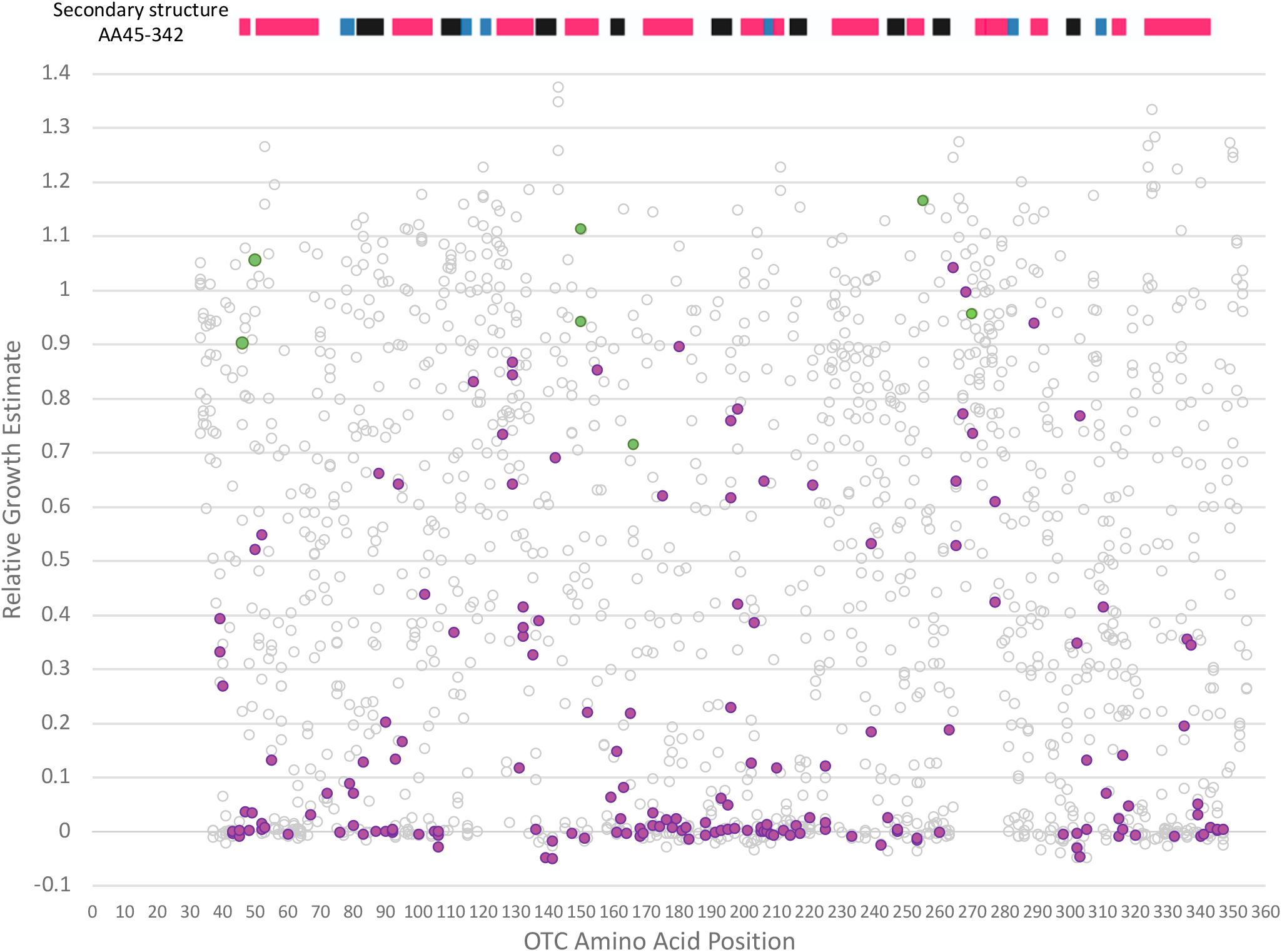
Relative growth of amino acid substitutions versus position in yOTC. Shown in grey spots are the relative growth values of each yOTC substitution. Shown in purple are amino acid substitutions corresponding to ClinVar pathogenic, pathogenic/likely pathogenic, and likely pathogenic variants. Shown in green are amino acid substitutions corresponding to ClinVar benign, benign /likely benign, and likely benign variants. Protein secondary structure is also depicted above for amino acids 45-342 (alpha helix – pink, turn – blue, beta sheet – black).

While OTC shows a general intolerance to amino acid substitutions, we expected that altering the amino acid at an active site of the enzyme would have a particularly high likelihood of impairing OTC function and therefore inhibiting growth in our assay. OTC has several known active sites involved in recognition and binding of carbamoyl phosphate and ornithine for enzymatic conversion to citrulline. Binding of carbamoyl phosphate involves the SxRT (S90, T91, R92, T93) and HPxQ (H168, P169, I170, Q171) motifs as well as residue R141. Most amino acid substitutions at these positions in our library displayed null or low levels of activity (Supplemental Table S3). The D263 residue of the DxxxSMG motif (D263, S267, M268, G269), which participates in the binding of ornithine, is highly intolerant to changes as all substitutions tested displayed null or low levels of activity. The HCLP (H302, C303, L304, P305) motif participates in binding both substrates, and the majority of changes in this motif also displayed null or low levels of activity. Taken together, amino acid substitutions at these active sites are significantly enriched (1-sided Fisher’s Exact Test; p-value = 1.9 × 10^−8^) for null levels of activity (relative growth <5%), relative to the rest of the protein.

Finally, we evaluated the agreement of our results with expected representation in the human population. Because of the severity of this X-linked disease, we expect that *OTC* missense SNPs that result in deleterious amino acid substitutions will be underrepresented in the human population. Consistent with this, amino acid substitutions corresponding to human missense SNPs present in the gnomAD database^36^ have significantly higher (p<0.001) median growth in our assay than random samples of the same size drawn from the full set of amino acid substitutions. Thus, although the proportion of deleterious amino acid substitutions is relatively high, our results agree with expectations based on the known features of the protein structure and their representation in the human population.

### Assay Results Agree Closely with ClinVar Clinical Annotation

To evaluate how our functional assay compared with human phenotypes, we assessed the growth scores of amino acid substitutions corresponding to those caused by missense SNPs with pathogenic and benign calls in the ClinVar database (Figure 6). Our library contained all amino acid substitutions corresponding to ClinVar benign SNPs (n= 3). Two of these (K46R and T150I) exhibited wild-type levels of growth in our assay. The remaining substitution (L166F) displayed a moderately high relative growth value of 71%. Our library contained four of the nine amino acid substitutions (G50A, T150N, H255R, and Q270R) corresponding to missense SNPs with ClinVar benign/likely benign and likely benign classifications. All four of these exhibited relative growth levels near or above wildtype yOTC activity. Thus, although the number is relatively small, our assay results agree well with the benign and likely benign clinical significance annotations of variants currently in ClinVar.

**Figure 6.**
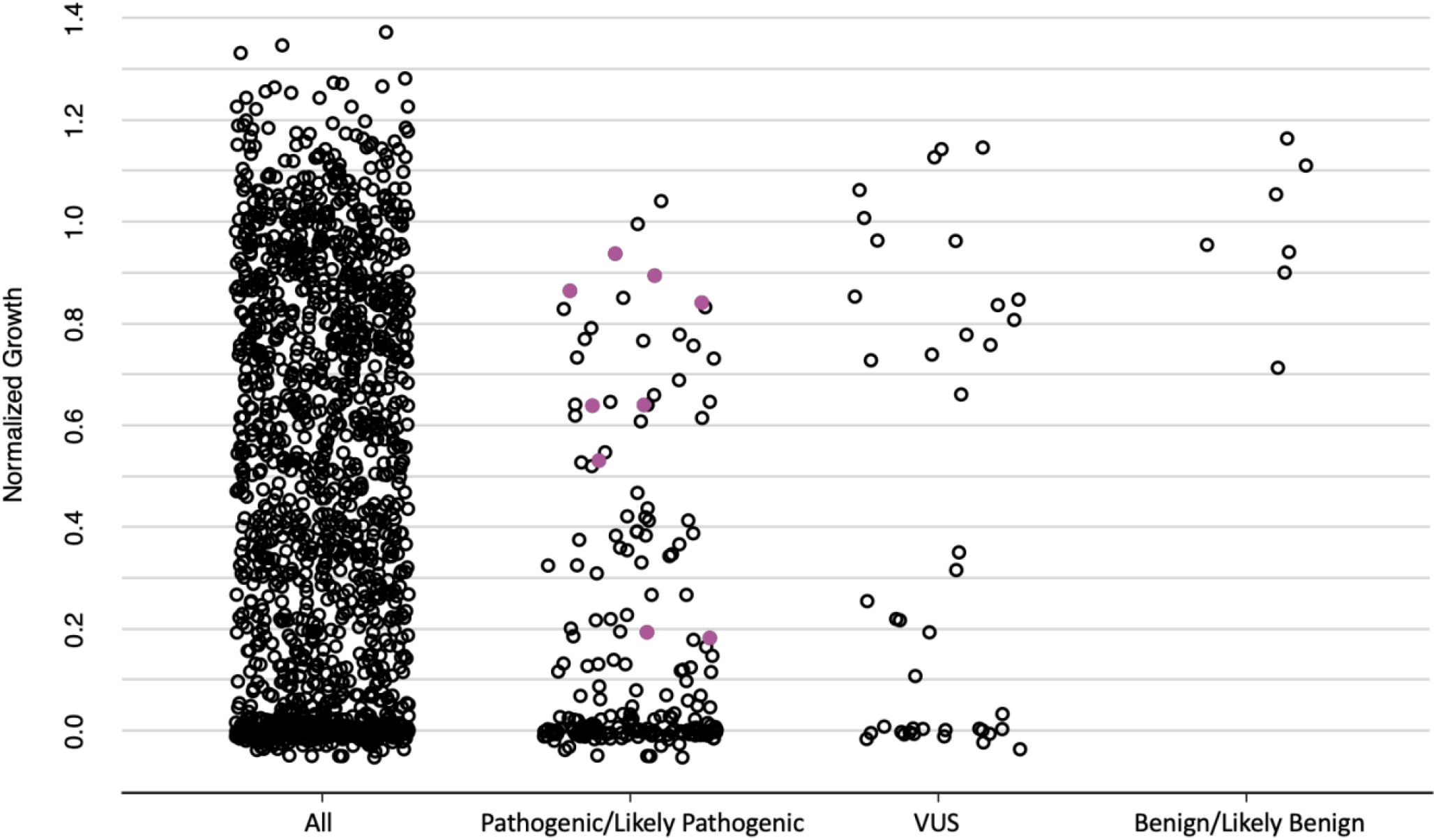
Strip charts of normalized growth for all amino acid substitutions and those corresponding to ClinVar classification groups. Shown in purple are amino acid substitutions corresponding to missense SNPs predicted to impair splicing.

We next analyzed the set of amino acid substitutions in our library corresponding to SNPs with ClinVar pathogenic (n= 165 of 208) or pathogenic/likely pathogenic and likely pathogenic (n= 30 of 39) classifications. The majority (~57%) of both groups fell below the stringent high-confidence cutoff of less than 5% relative growth that defines the null activity range of our assay (Figure 7). In contrast, only 27% of all assayed amino acid substitutions have growth in this range. Thus, the enrichment of amino acid substitutions corresponding to SNPs with pathogenic/likely pathogenic classification in the null range (relative to all other substitutions) is highly significant (1-sided Fisher’s Exact Test; p-value = 1.2 × 10^−20^). In addition, after excluding all amino acid substitutions in the null range, the remaining amino acids associated with pathogenicity still have significantly reduced median growth relative to random samples of the same size (p<1×10^−4^).

**Figure 7.**
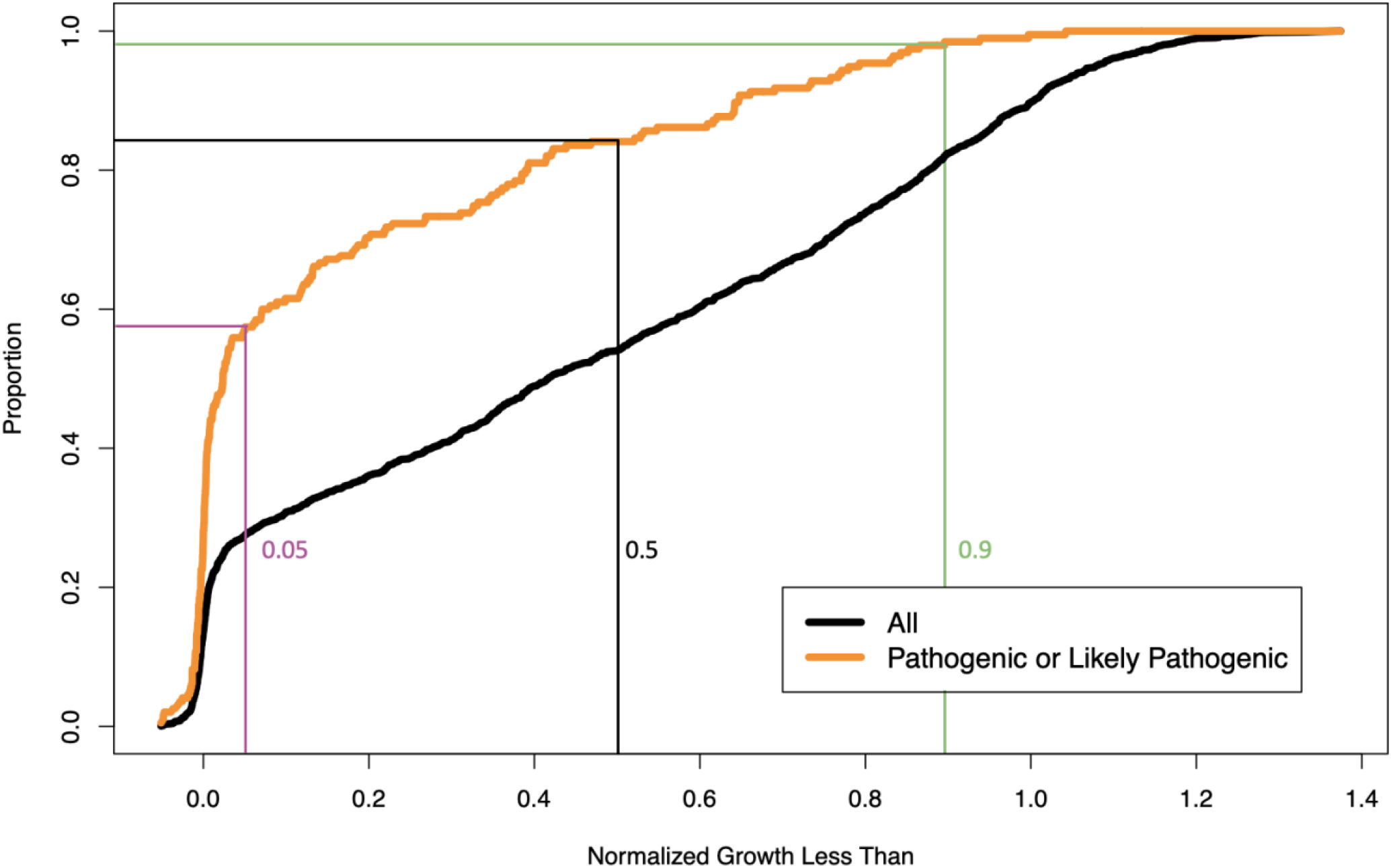
Comparison of the proportion of all amino acid substitutions and those with ClinVar pathogenic/likely pathogenic annotation falling under a given level of normalized growth.

The phenotypic distribution of the benign-associated and pathogenic-associated amino acid substitutions are well separated in our assay with only a small degree of overlap between them. For amino acid substitutions corresponding to pathogenic and likely-pathogenic variants, only 8% (n= 14 of 165) and 7% (n= 2 of 30) respectively, fall above the 71% relative growth value of the worst-growing benign-associated amino acid substitution. In comparison, 33% of all amino acid substitutions have growth in this range. Taken together, the strong correlation between the clinical significance calls in ClinVar and the quantitative scale of our assay supports the validity of the yeast assay for assessing the human protein function. In addition, with the caveat that the number of amino acid substitutions associated with benign annotations is small, the degree of separation between the two distributions suggests that a relatively clear classification threshold between the two clinical classes may exist using the growth scale of our assay.

### Variant Assay Values Agree Closely with OTC Clinical Stratification

In males, OTC disease presentation is classified into two groups based on age of onset, neonatal and late-onset (>6 weeks). Amino acid substitutions corresponding to SNPs with neonatal onset are expected to more severely impair OTC activity than those corresponding to the late-onset alleles. It has been suggested that disease presentation in females results from skewed patterns of X-inactivation affecting expression of a wild type and a severe pathogenic SNP.^37–39^ We therefore examined the behavior in our assay of amino acid substitutions corresponding to SNPs falling into these three groups (Figure 8).

**Figure 8.**
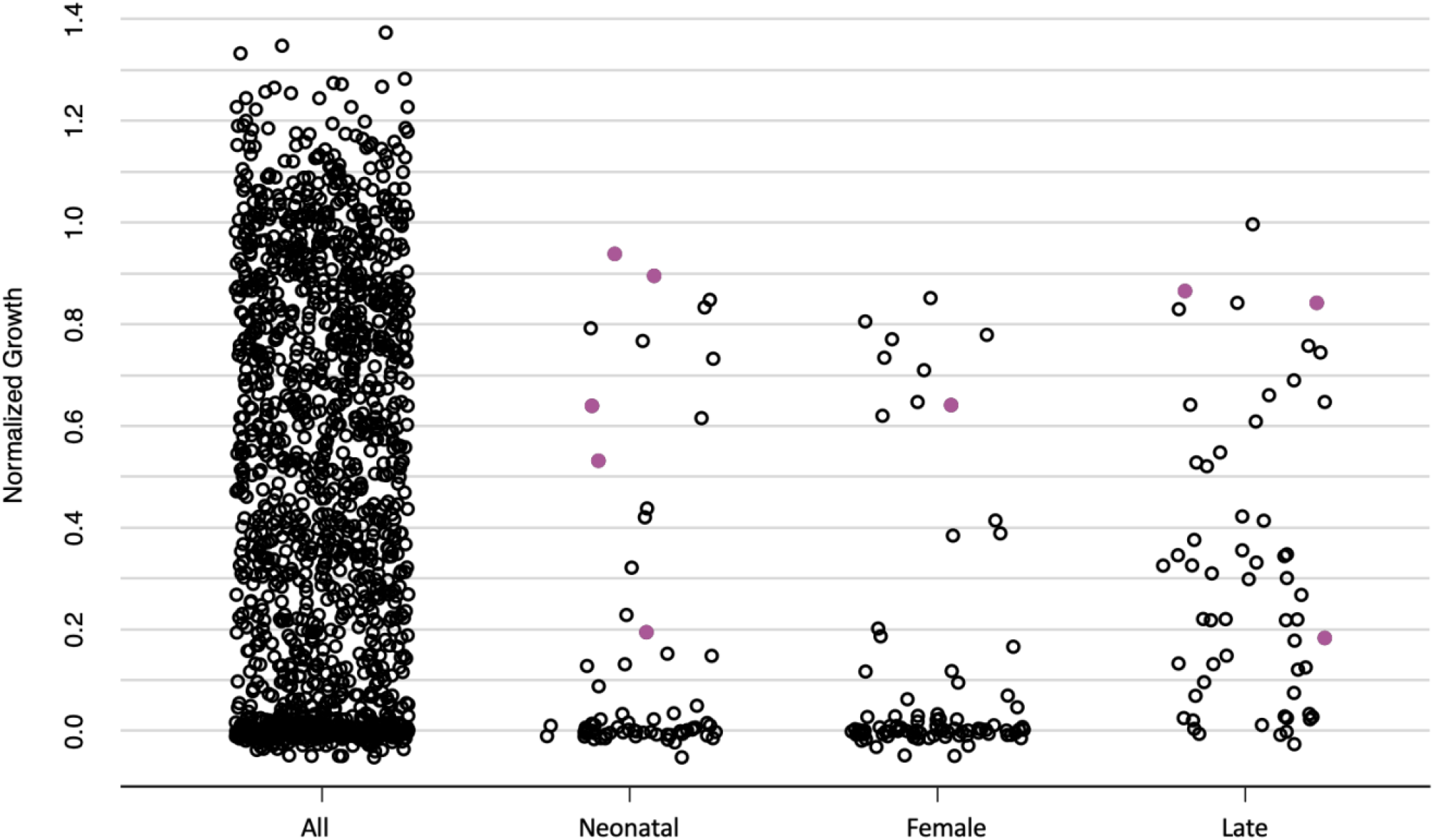
Strip charts of normalized growth for all amino acid substitutions and those corresponding to UCDC classification groups. Shown in purple are amino acid substitutions corresponding to missense SNPs predicted to impair splicing.

The distributions of amino acid substitutions corresponding to neonatal and female variants were very similar and associated with severe loss of activity in our assay (Figure 9). Of the 69 neonatal-associated substitutions present in our library (n= 69 of 91), 70% had activity in the null range of our assay (<5% relative growth). Similarly, of the 88 female-associated substitutions in our library (n= 88 of 107), 77% had growth values below the 5% cutoff. In contrast, only 27% of all amino acid substitutions fell in this growth range (Figure 9). These results support the hypothesis that female presentation, like neonatal presentation in males, is more closely associated with severe loss of function variants. Relative to all other amino acid substitutions, this enrichment of neonatal- or female-associated substitutions in the null range was highly significant (1-sided Fisher’s Exact Test; p-value = 9.8 × 10^−38^). Further, after excluding all amino acid substitutions in the null range, the combined set of the remaining neonatal- and female-associated substitutions still had significantly lower median growth relative to random samples of the same size (p<0.01). In particular, amino acid substitutions in these two classes were almost completely absent from the set of strains that showed growth similar to wild type (defined here as >90% relative growth), which make up 18% of all amino acid substitutions (Figure 9). Thus, amino acid variants causing complete or near-complete loss of OTC function in our assay are strongly associated with neonatal and female presentation of disease.

**Figure 9.**
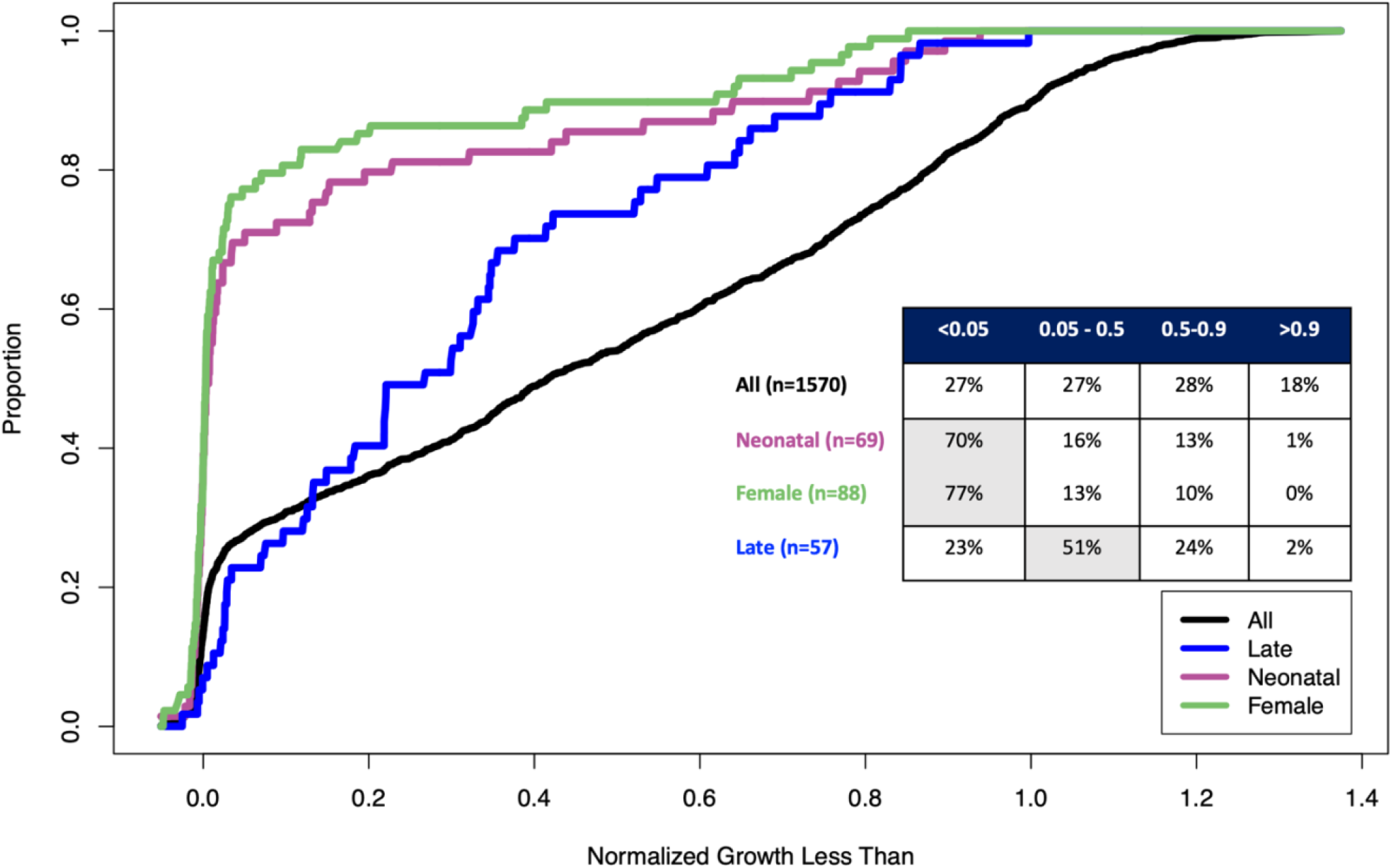
Comparison of the proportion of all amino acid substitutions falling under a given amount of normalized growth versus those associated with neonatal, female, or late presentation.

Amino acids substitutions in our library that correspond to late-onset variants (n = 57 of 71) show significantly lower median growth relative to random samples of the same size drawn from all amino acid substitutions not assigned neonatal or female classifications (p<0.01). In particular, of the 57 late-onset variants in the dataset, only one was observed among strains with growth similar to wild type (>90% relative growth). Therefore, substitutions corresponding to late-onset variants exhibit impaired growth in our assay, as expected for amino acid changes encoded by pathogenic alleles. However, the median relative growth of late-onset associated substitutions is shifted significantly upward from that of neonatal and female associated substitutions (p <1 × 10^−5^ using the empirical null distribution of median differences). While approximately 75% of substitutions with neonatal or female classifications reside in the null range of the assay, most of the late-onset associated substitutions (51%) had growth values between 5% and 50%, with the remainder split evenly between the <5% and 50-90% growth ranges (Figure 9). Therefore, late-onset presentation is associated with amino acid substitutions that have impaired activity in our assay, but generally less impaired than the amino acid substitutions associated with neonatal and female presentation.

Together, these results demonstrate a very close agreement between growth in our assay and disease stratification. All three classes of disease-associated amino acid substitutions are depleted in the upper range of our assay, but those associated with neonatal and female presentation show complete, or near complete, loss of OTC function, while those associated with the late-onset class demonstrate less severe loss of OTC function.

### Heterogeneity in Age of Diagnosis in the Late-Onset Class

The correlation between age of disease onset and activity in our assay is strong. Amino acid changes corresponding to neonatal variants (<6 weeks old onset) display significantly lower median growth than those corresponding to late-onset variants (>6 weeks old onset). However, within the late-onset class, the age of diagnosis varies greatly, from 6 weeks of age up to the 7^th^ decade of life (Figure 10). Therefore, we extended our analysis to investigate whether there was also a correlation between age of (late) onset and growth in our assay. To this end, we further resolved the exact age of onset or diagnosis in the late-onset class of males by performing an exhaustive literature search for these parameters (Supplemental Table S6). Using these data, we observed a significant negative correlation (p<3.7 × 10^−7^) between age of onset in affected males and growth of the corresponding amino acid substitutions (Figure 11). However, this relationship only explained a small proportion of the variance in age of onset (r^2^=0.21). In addition, our analysis highlighted a number of instances where groups of patients with the same amino acid substitution (A208T, P225T, R40H, V337L, R277W) displayed a wide range of age of onset (Supplemental Figure 1). Together, these results indicate that, while late-onset variants are associated with moderate loss of function in OTC, the actual age of onset within the late class is only weakly determined by the relative severity of the OTC allele. This is consistent with the known role of environmental triggers among late-onset cases of hyperammonemia (Table S6).

**Figure 10.**
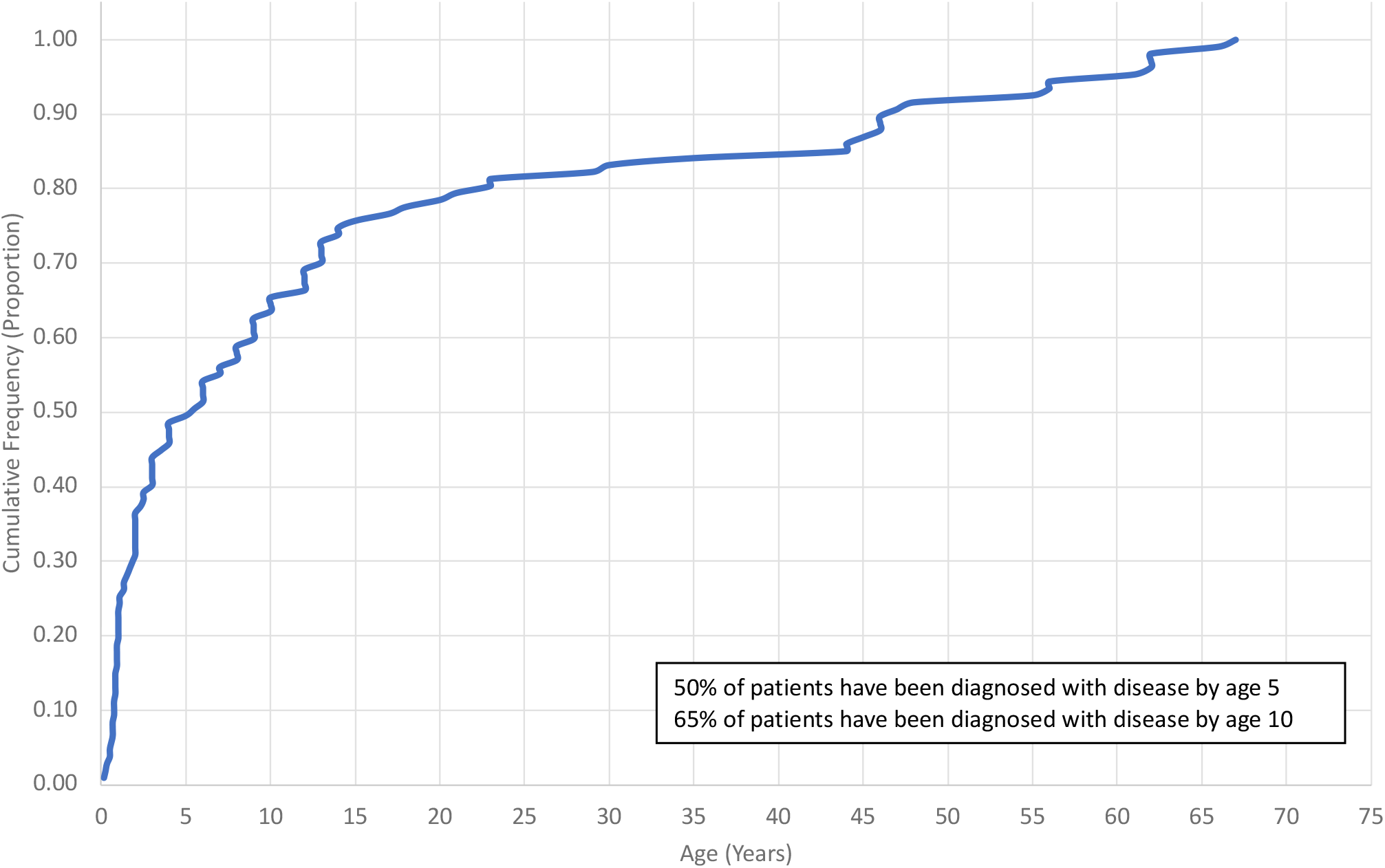
Literature curation: age of onset/diagnosis for symptomatic, late onset males.

**Figure 11.**
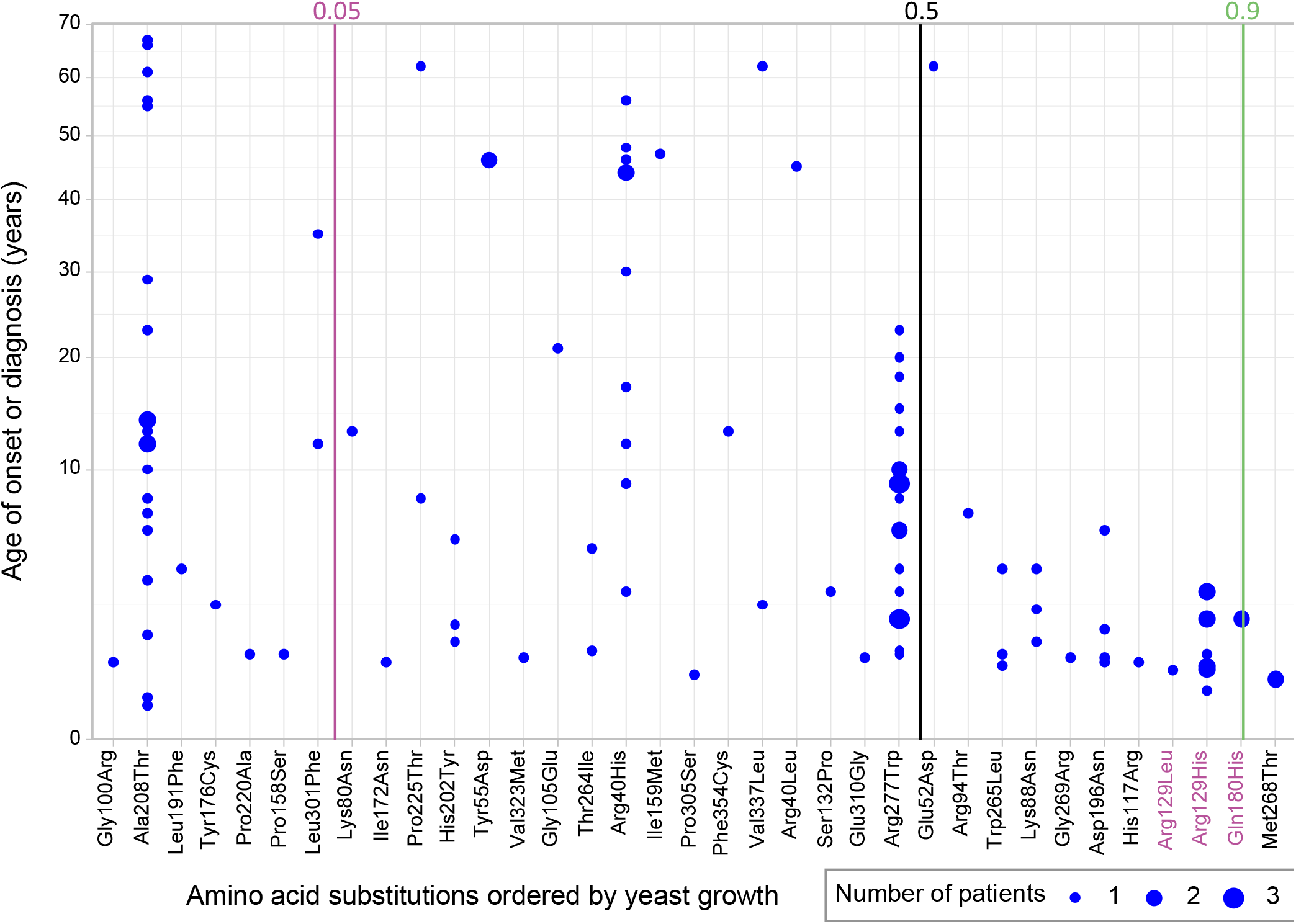
Age of onset of late male patients and yeast growth scores for accompanying genotypes. Labeled in purple are amino acid substitutions corresponding to missense SNPs predicted to impair splicing.

### Separating the Effects on Splicing of Missense Variants from the Effects on Amino Acid Sequence

Missense mutations affecting the first and last codons in an exon can affect splicing in addition to causing amino acid substitutions. Both effects can impact protein function and can be difficult to separate. Because our assay measures the effect of amino acid substitutions and uses a construct (*yOTC*) that lacks introns, the effect of a variant on the amino acid sequence can be assessed independent of any potential effects it might have on the function of splice donor and acceptor sites.

On this basis, we analyzed all possible missense SNPs in the last 3 bp of *OTC* exons 2-9, and the first bp of exons 3-10 for their predicted effect on splicing. The junction between exons 1-2 was not assessed since exon 1 encodes the first 26 amino acids that are a part of the *OTC* mitochondrial leader sequence, which is not present in our expression construct. This identified a set of 76 SNPs on which we performed *in silico* predictions for their effect on OTC mRNA splicing using SpliceAI.^40^ These 76 SNPs correspond to 68 unique amino acid changes, of which 58 were present and measured in our assay (Supplemental Table S4).

Within the set of 58 amino acid substitutions, we have growth measurements for thirteen that are associated with variants having ClinVar pathogenic or likely pathogenic classifications. We compared the predicted effect on splicing for these variants to the results of our growth assay in order to resolve the likely mechanism of pathogenicity. Four of these thirteen (R129H, R129L, Q180H, K289N) have a SpliceAI Δ score ≥ 0.83 (Δ score range of 0-1 representing the probability that a variant will negatively affect splicing) but a high growth value in our assay (>84% growth). This suggests that the amino acid substitutions associated with these SNPs have little effect on OTC function, and instead, they are pathogenic because they cause splicing defects. Notably these four SNPs encode 4/9 of all pathogenic variants associated with amino acid substitutions having growth >80% in our assay (Figure 6).

Among the remaining nine pathogenic SNPs, four have a SpliceAI Δ scores **≤** 0.5 and display low growth (<36%) in our assay. These variants are unlikely to cause splicing defects but the amino acid substitutions that they encode strongly impair OTC function, suggesting this is the mechanism of pathogenicity. The final five SNPs have both a SpliceAI Δ score ≥ 0.79 and display impaired growth (<65%) in our assay. Therefore, these SNPs both impair splicing and encode amino acid substitutions that have a deleterious effect on OTC function.

Thus, we can effectively discern if the pathogenicity of a variant is likely due to deficits in proper splicing alone, low enzymatic activity alone, or both improper slicing and low activity, in which case any protein translated from the reduced proportion of properly spliced mRNA is still subject to lower enzymatic activity.

### Defining Growth Thresholds for Functional Annotation of Variants

A major goal of our study was to use experimental evidence from our assay as a criterion for the reclassification of Variants of Uncertain Significance (VUS). In order to convert our assay results into functional evidence for impairment or lack of impairment of OTC activity, we need to identify growth thresholds based on the behavior of variants with known pathogenic and benign clinical interpretations. We can then leverage our results for this set of clinically characterized variants to make inferences about the large number of remaining, clinically uncharacterized variants from our comprehensive dataset.

First, we set out to identify a growth threshold below which OTC activity is impaired to a level consistent with known pathogenic variants. Below this threshold, we expect to see enrichment for existing pathogenic variants and depletion for benign variants. The lowest-growing amino acid substitution corresponding to a benign missense variant shows 71% growth in our assay, and only 8% of pathogenic or likely pathogenic, variants are associated with substitutions having growth higher than this (versus 33% of all substitutions) (Figures 6 & 7). Notably, several of these outlier variants are predicted to impair splicing (Figure 6), suggesting that this, rather than their effect on protein sequence, explains their pathogenicity. Therefore, we believe that growth less than 71% in our assay provides strong evidence for functional impairment of OTC to a level consistent with existing pathogenic variants. This cutoff indicates that a very high proportion (1049/1570, 67%) of all SNP-accessible OTC amino acid substitutions cause functional impairment strong enough to be potentially pathogenic.

In addition, this functionally impaired range can be further subdivided, with <5% growth vs 5-71% growth in our assay associated with differences in disease presentation. Amino acid substitutions with growth <5% in our assay form an approximately normal distribution around the value of the null control strain (defined to have normalized growth of zero), consistent with this class representing total loss of function alleles. Amino acid substitutions associated with neonatal variants are very strongly enriched in this class (Figures 8 & 9). In contrast, amino acid substitutions associated with late-onset variants are most enriched in the lower range of our assay, but above 5% activity, i.e. moderate but not complete loss of OTC function.

Next, we set out to identify a threshold identifying a minimum level of OTC activity consistent with known benign variants. Above this threshold, we expect depletion for known pathogenic variants and enrichment for known benign variants. Amino acid substitutions corresponding to pathogenic and likely-pathogenic variants are depleted for growth >90%, with only 3/194 falling in this range (versus 18% of all amino acid substitutions). There are only a few existing OTC variants with benign or likely benign clinical classifications, but all but one of these in our library has growth >90% in our assay (Figures 6 & 7). Therefore, human missense variants corresponding to amino acid substitutions in our assay with >90% growth result in levels of OTC activity consistent with existing benign alleles, i.e. not functionally impaired. However, these missense variants could still be pathogenic if they have an additional deleterious effect beyond the associated amino acid substitution, e.g. on splicing.

The growth range 71-90% in our assay remains ambiguous in terms of functional evidence for OTC activity. This range is higher than the lowest growing benign-associated amino acid substitution but is lower than several pathogenic-associated substitutions (Figure 6). Additional clinical information is likely to help further resolve this range.

### Reclassifying VUS Using Functional Assay Data

Having identified functional thresholds in our data, we set out to use our results to carry out OTC variant reclassification. We have identified growth <71% in our assay as functional impairment of OTC, but for our reclassification we used the highly conservative 5% threshold associated with complete loss of OTC function. Although the 5% cutoff is very stringent, it captures 57% of amino acid substitutions corresponding to ClinVar pathogenic and likely pathogenic variants as well as 70% of those corresponding to neonatal variants and 81% of those corresponding to variants present in affected females. In total, 27% of all amino acid substitutions that we tested had growth <5%, including 320 corresponding to missense variants that have yet to be documented in the human population. Treating growth <5% in our assay as strong functional evidence for impaired OTC activity, we examined the effect this evidence would have on variant classification using the current ACMG/AMP guidelines.^41^

Of the 39 amino acid substitutions in our library that correspond to variants annotated as VUS in ClinVar (n= 39 of 58), 16 had growth values below our stringent 5% cutoff (Figure 6). Additionally, 17 likely-pathogenic variants and 1 “conflicting interpretations of pathogenicity” variant had corresponding amino acid substitutions with growth values below 5%. We used current ACMG/AMP guidelines to reanalyze these 34 variants (Table 1 & Supplemental Table S5). Without our data, 3 were classified as likely-pathogenic and 31 as VUS. However, inclusion of the yeast assay data as PS3/BS3 functional evidence reclassified 21 of these VUS to likely-pathogenic. The classification of the 3 likely-pathogenic variants did not change with the inclusion of our yeast data.

**Table 1.** Reclassification of VUS, likely pathogenic, and conflicting variants according to current ACMG/AMP guidelines.

|  | ClinVar | ACMG<br>alone | ACMG with<br>yeast data |
| --- | --- | --- | --- |
| **VUS** | 16 | 31 | 10 |
| **Likely Pathogenic** | 17 | 3 | 24 |
| **Conflicting** | 1 | 0 |  |

## Discussion

Technological advances are enabling the integration of genome sequencing into routine healthcare, including proposals and large pilot programs to study the impact of including DNA sequence-based diagnostics in newborn screening.^42^ While many human genes are clinically actionable, i.e. the presence of a pathogenic variant changes medical management, currently only ~2% of germline missense variants have clinical interpretations.^43,44^ The remaining variants of uncertain significance offer no information to inform diagnosis or guide treatment. The scale of the problem is immense and is a major obstacle for realizing the goals of precision medicine. As such, eliminating the VUS category of clinical diagnosis by 2030 has been deemed one of the ten “highest-priority elements envisioned for the cutting-edge of human genomics”.^45^ Because computational prediction algorithms often give inaccurate or conflicting results,^46^ large-scale functional assays are the only means of variant interpretation currently poised to match the pace of variant discovery.

Here, we describe the development and application at scale of such an assay, a yeast-based quantitative growth assay measuring the activity of the human OTC protein. OTC Deficiency is a devastating disease for which rapid detection in the newborn period is often needed to significantly reduce morbidity and mortality. As the OTC gene is located on the × chromosome, the majority of patients affected with OTCD are hemizygous males. Because our assay is conducted with single copy gene variants integrated into a haploid yeast strain, our functional assay not only assesses the activity of individual OTC variants, but also recapitulates the genotypes of male patients with single amino acid changes in the enzyme encoded by this X-linked gene. In total, we measured the effect of 1,570 single amino acid substitutions on OTC activity and determined that 1,049 had levels of functional impairment consistent with known pathogenic variants.

The validity of the assay is supported by the close agreement with existing information about OTC, both at the level of clinically characterized variants and protein structure. Amino acid substitutions corresponding to known pathogenic variants and substitutions affecting active sites in the protein both demonstrated impaired activity in our assay. In addition, the degree of impairment shows good agreement with clinical stratification of known disease variants. Amino acid substitutions corresponding to the most severe clinical variants (neonatal presentation in males) are highly enriched in the null range of the assay. Additionally, our results support the long-standing hypothesis that female presentation occurs in those harboring highly deleterious variants. Amino acid substitutions corresponding to male late-onset OTCD variants also display significant impairment in our assay, but on average have activity levels that are significantly higher than substitutions associated with severe variants. Interestingly, the relatively weak correlation between these more moderately impaired variants and finer scaled analysis of age of onset, is consistent with observations that there is a strong environmental component to late-onset presentation. In these cases, clear triggering events are often identified. These events include dietary changes, surgeries, infections, pregnancy, or exposure to valproate (which inhibits the upstream NAGS enzyme in the urea cycle) or glucocorticoids (which inhibit protein synthesis and promote catabolism). As these events are stochastic, age of late-onset for moderate variants is also largely stochastic. This pattern is consistent with moderately impaired OTC alleles being sufficient for baseline urea cycle activity, but not sufficient under stress conditions. Thus, knowledge of variants with moderately impaired growth values (5-71%) are also highly actionable as patients can avoid potential hyperammonemia triggering events and physicians can more rapidly diagnose metabolic crises should they occur.

