## Supplemental File 1 for "The functional impact of 1,570 SNP-accessible missense variants in human *OTC*"

**Determining OTC Variant Encoded by Each Isolate Using MinION Sequencing**

Because each yeast transformation was carried out with a mixture of all SNP-accessible variants at a specific target codon, we know the codon changed in each transformant, but not which variant it was transformed to. To determine this, and to identify any secondary mutations, we developed a pipeline utilizing Oxford Nanopore MinION sequencing.

First, the 96-well variant plates (1 variant per well) were collapsed in sets of 16, so that well A1 on each collapsed plate combined the variants in well A1 in each of 16 variant plates. Variant plates were ordered so that 16 different codons were targeted in each collapsed well. For each collapsed well, we know the 16 target codons and we also know the set of potential variant sequences introduced at each codon by Twist Biosciences. By sequencing pooled DNA in each well of each collapsed plate and identifying the most common expected variant sequence at each target codon, we can determine the identity of the variant in each well of the variant plates.

After collapsing the plates, fragments encompassing the *ARG3* promoter, *yOTC* ORF, and *TRP2* terminator were amplified from the pooled genomic DNA in each collapsed well using 16 or 18 cycles of PCR with dual barcoded primers (identifying each well). ONP adapters were added on via ligation. Sequencing was performed on an Oxford Nanopore MinION Mk1B with a Flongle adaptor & flowcell with 1 collapsed plate per run.

After demultiplexing, reads were aligned to the *yOTC* reference sequence in R (R Core Team (2022). R: A language and environment for statistical computing. R Foundation for Statistical Computing, Vienna, Austria. URL https://www.R-project.org/) using the pairwiseAlignment function of the Biostrings package (Pagès H, Aboyoun P, Gentleman R, DebRoy S (2022). Biostrings: Efficient manipulation of biological strings. R package version 2.64.1, https://bioconductor.org/packages/Biostrings) using the with parameters gapOpening=10 and gapExtension=2. For each read, alignments were carried out in both orientations and the highest scoring alignment was kept. Then, for each of the 16 target codons in each well, the most frequent and second most frequent potential Twist variant codons were recorded. The most frequent variant was provisionally accepted if the ratio of first variant codon counts to reference codon counts was >0.02, based on the multiplexing expectation of 1/15 (0.061) along with some sampling variation. The set of reads containing the variant codon were then scanned for secondary mutations affecting >50% of the reads, which were then recorded. Secondaries that were not indels were classified as synonymous, missense or nonsense.

The observed frequency of each provision variant codon was then compared to the frequency expected under a variant-specific error model. To generate these models, for each well we recorded the counts of all potential Twist variants at codons NOT among the set of 16 target codons in the well. In addition, we recorded the counts of the reference codon at these positions. These data were combined to estimate an error frequency for each potential variant codon. Provisional variant calls were accepted if their observed frequency was >3.3 times their error frequency. In addition, we required that this ratio was higher than the equivalent ratio for the second most frequent variant.

**Validating the MinION Variant Calls Using Illumina Sequencing**

To validate our Oxford Nanopore pipeline, we performed amplicon sequencing on the Illumina sequencing platform to confirm the variant in each well of each variant plate. These calls were then compared to the result from the Oxford Nanopore pipeline. For the Illumina sequencing, the *yOTC* ORF was divided up into four ~300bp overlapping segments. The specific segment to be amplified and sequenced corresponded to the known codon position of the variant within our library. 35 cycles of PCR amplification were performed using genomic DNA as a template. The initial PCR reaction was then diluted and used for an additional 24 cycles of PCR to add on Illumina index sequences. The final PCR reactions were pooled and gel purified before performing Illumina sequencing on a Nextseq 500 using 300-cycle, dual-indexed, paired-end sequencing. Reads were demultiplexed and aligned to the *yOTC* reference sequence as described.[1] For each isolate, the basecalls across the target codon position were used to identify the variant codon sequence.

Comparison of the MinION and Illumina results revealed almost complete agreement (99.8%) for the MinION calls passing the quality filters (Supplemental Figure 1). The great majority of the remaining "variants" are occasions when, as determined by the Illumina sequencing, no change had actually been made at the target codon (i.e. wild type *yOTC* sequence). As expected, for these "no-change" isolates, the most frequent potential variant identified by the MinION pipeline occurred at a low frequency similar to the error frequency for that variant (Supplemental Figure 1). In contrast, when a variant was present, it was observed at the frequency expected from the level of MinIon multiplexing (16-fold) and was correctly identified with high accuracy.

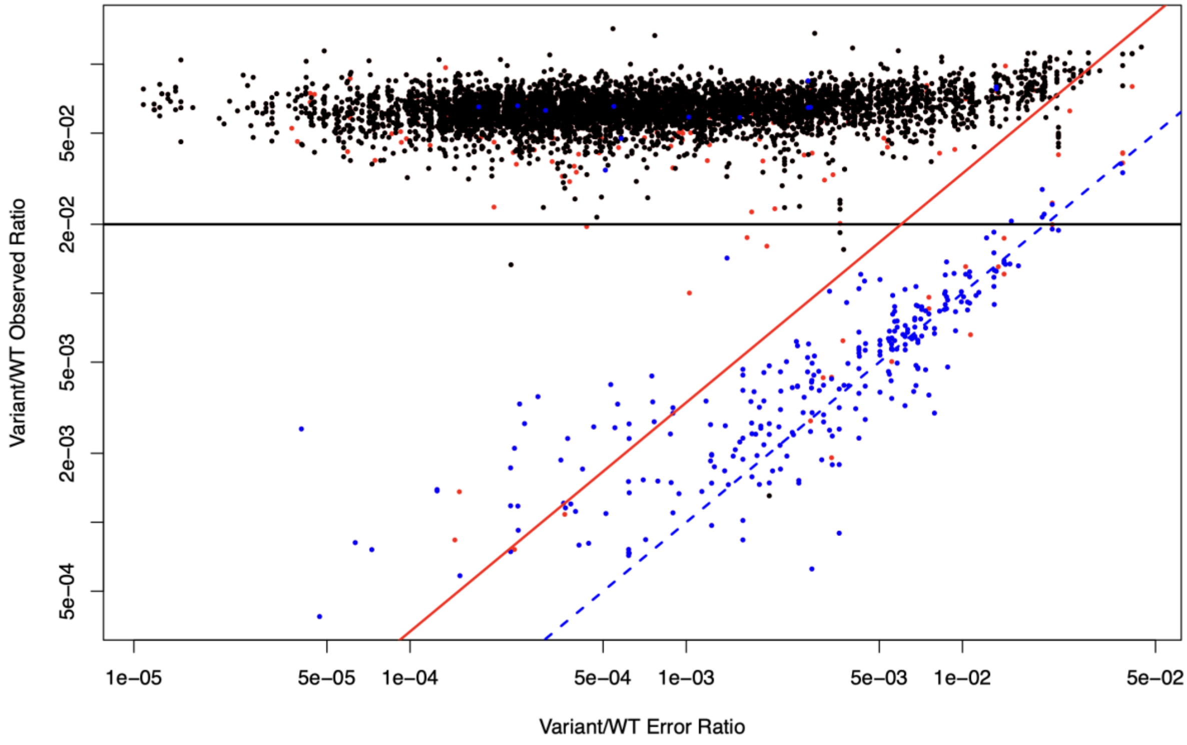

**Supplemental Figure 1. Agreement between *yOTC* variant codons identified by Oxford Nanopore versus Illumina sequencing.** For each variant call, the observed ratio of Oxford Nanopore variant to reference reads (expected to be 1/15) is plotted against the empirically calculated error ratio. Blue dotted line indicates 1:1 relationship between observed and error ratios. Variant calls in agreement with Illumina shown in black. Disagreements where Illumina identifies no change at the target codon (i.e. reference sequence) shown in blue. Remaining disagreements shown in red. Filters require observed variant to reference ratio to be above 0.02 (black line) and to be >3.3 times error rate (red line).

### Growth Assays and Imaging

Yeast transformation isolates were picked and grown in rich medium supplemented with drug selection and then pinned onto solid rich medium containing glycerol as the carbon source (YPG), to eliminate any petite yeast cells (lacking mitochondria). Cells were then pinned from YPG solid medium back into nonselective rich liquid medium (YPD) and grown to saturation overnight at 30° C. Cultures were then pinned onto solid omniplates containing minimal (SD) media (no arginine) using a Biomek i7 robot outfitted with a V&P 96-pin head. Source plates were pinned onto SD plates in triplicate and grown at 30C for 72 hours, with photographs being taken every 24 hours. Plates were photographed with a mounted Canon PowerShot SX10 IS compact digital camera under consistent lighting, camera to subject distance, and zoom. Images (ISO200, f4.5, 1/40 second exposure) were acquired as jpg files.

### Phenotyping Image Analysis

The area (in pixels) and intensity (gray scale value) of each replica-pinned patch of cells was extracted from each image with a custom in-house script. Phenotype values were extracted from plate images using a custom pipeline developed in python, primarily using the sci-kit image library as follows. First, each omniplate image was cropped into 96 square tiles, each tile centered at a pin spot location so that tiles contain a single patch of cells. Identical camera set up and nearly identical plate positioning during plate imaging allowed a single set of reference locations to be used for this preprocessing step. An additional preprocessing step was performed to mask any bright objects, such as reflections from the agar meniscus or parts of the omniplate edge. This consisted of converting the 8-bit color images to grayscale, intensity thresholding with a value automatically selected via Otsu's method and disregarding any resulting objects that intersect with the tile edge in downstream processing.

After preprocessing, the pixels constituting the patch were identified in the grayscale tiles using either intensity thresholding or circle detection. First, Otsu's method was used to automatically select a threshold value. If the threshold returned from Otsu's method, $\tau$, was sufficiently different from the average background agar intensity ($\tau > 0.55$), it was applied to the grayscale image to create a binary image. However, if the Otsu threshold was similar to the background agar intensity, as in the cases of null growth, then the threshold was increased slightly ($\tau$ ß $\tau + 0.01$) to improve the separation between patch and background intensities. Using a higher threshold value increases the chance that binarizing will distinguish the patch from the agar background. Next, the morphological operations of filling and closing were applied to the binarized image produced by thresholding. The filling function identified objects, from which the largest one was selected, and the closing function ensured that the identified patch region did not have any unselected pixels within its boundary. If this returned a sufficiently large object (> 2000 pixels) contained completely within the tile, it was assumed to be the cell patch and this object was used as the tile mask.

If no sufficiently large object was detected via thresholding, then a mask identifying the patch was produced with circle detection instead. Circle detection was performed by binarizing the tile with an adjusted mean threshold ($\tau$ = mean(grayscale tile) + 0.02) and applying the Circular Hough transform with a minimum radius equal to the initial pin radius (25 pixels). This exploits the geometry of the pinned cells to capture any pixels within patches where growth is too faint to be detected with intensity thresholding. The most probable circle identified by the transform was used as the patch mask, provided it did not intersect the edge of the tile.

Once a mask identifying which pixels belong to the patch vs the background was produced for each tile, the grayscale tile was segmented by applying the mask to the grayscale image. At this point an additional visual check was performed by overlaying detected patch masks on the original images and displaying them. For a small number of images, faintly growing patches in corner positions were not detected so these plates were reprocessed with different reference points for the initial cropping step. After segmentation, the final patch area is simply the total number of pixels in the patch and the patch volume ("IntDen") is the sum of the grayscale values of these pixels. Note that the parameter values used here, such as minimum pixel radius and minimum patch area, are specific to the camera settings and imaging set up used in this experiment. The code used to process images is available at https://github.com/lacyk3/pnri-projects/tree/Image-Analysis-Demos/OTC%20Project. The functions used for preprocessing and segmentation have also been further developed and the most recent version can be downloaded from https://github.com/lacyk3/PyPl8.

**Phenotype Normalization and Quality Control**

The raw growth phenotypes underwent a series of normalization and quality control steps, carried out using a custom script. Before any other steps, to adjust for pin size variation, phenotype estimates were multiplied by pin-specific normalization factors, estimated using control plates made up entirely of wild-type strains grown for 24 hours. Remaining normalization steps were based on the two null, and four wild type, control patches present on each plate.

We expect that the remaining systematic effects on growth will not affect null controls or isolates with null-like growth and will scale with growth level (above null). Therefore, these effects were modeled as multiplicative, after first subtracting the mean value of the null-control patches across all plates.

First, the three repeat pinnings of each isolating plate were normalized. A coefficient for each repeat plate was calculated so as to make the mean wild type control growth value on each plate equal to the mean wild type across all repeats of that plate. This coefficient was then applied to all growth estimates on the repeat plates.

Next, adjustments were made for plate-to-plate variation between variant plates and for the effect of mean neighbor patch growth. Neighbor patch growth has a significant effect on growth, as patches that are surrounded by poorly growing genotypes experience a boost in growth. A linear model with terms for plate and mean neighbor growth was then fit to the growth values of the wild type control strains. The coefficients from the model were then used to normalize all phenotype measurements.

At this point, all isolates that did not have a variant assignment that passed sequencing quality control, or which had any secondary indels, or secondary missense or nonsense mutations were removed. Isolates with multiple secondary mutations of any sort were also removed. Growth estimates for each variant and genotype (combining all isolates of that genotype) were then generated using linear models. Any isolate with an extreme estimated growth value, relative to its genotype, (greater than 500+1/0.85*Genotype_Estimate or less than -500 +0.85*Genotype_Estimate) was also removed.

Finally, normalization ratios for each edge were estimated, relative to the center of the plate. First, the mean growth value of each genotype was estimated for positions at each edge and for positions in the middle. When a particular genotype was missing from a region, an NA value was recorded. Then, using these values, linear models were used to estimate coefficients for each edge relative to the plate middle. These coefficients were then used to normalize all edge measurements.

#### Estimating the Relative Growth of Each Genotype

Finally, a linear model was used to estimate the normalized growth of each genotype. Values were then rescaled to set the null control as 0 and the wild type *yOTC* control as 1. Standard errors of growth relative to wild type were calculated as described.[1]
